# IHS: an integrative method for the identification of network hubs

**DOI:** 10.1101/2020.02.17.953430

**Authors:** Abbas Salavaty, Mirana Ramialison, Peter D Currie

## Abstract

Biological systems are composed of highly complex networks and decoding the functional significance of individual network components is critical for understanding healthy and diseased states. Several algorithms have been designed to identify the most influential regulatory points, or hub nodes, within a network. However, none of the current methods correct for inherent positional biases which limits their applicability. In addition, none of the currently available hub detection algorithms effectively combine network centrality measures together. To overcome this computational deficit, we undertook a statistical assessment of 200 real-world and simulated networks to decipher associations between centrality measures, and developed a novel algorithm termed “integrated hubness score” (IHS), which integrates the most important and commonly used network centrality measures, namely degree centrality, betweenness centrality and neighbourhood connectivity, in an unbiased way. When compared against the four most commonly used hub identification methods on four independent validated biological networks, the IHS algorithm outperformed all other assessed methods. Using this novel and universal method, researchers of any discipline can now identify the most influential network nodes.

## Introduction

The computational theory of complex systems, or graph theory, aims to provide a holistic, top-down, view of network interactions with the purpose of identifying critical network properties that reductionist approaches are incapable of identifying. Graph theory has been used for the investigation of complex networks within a broad variety of scientific fields including social networks, road traffic, telecommunications, cartography, chemistry, biochemistry, and biology in general (Balaban, 1985; Frainay & Jourdan, 2017; Hochberg & Ribó, 2018; Sandefur et al, 2013). In the age of high-throughput biological assays, systems biology techniques are being extensively used for the analysis of a variety of biological networks including gene regulatory networks (GRN), protein-protein interactions (PPI), and neural signals (Tieri et al, 2019). In these approaches, the topology of a network is analyzed and its different centrality measures (metrics demonstrating the local/global influence of each node within a network) are calculated to find deeper biological meanings and identify the most influential regulatory molecules, namely hub nodes. Hub nodes have high connections with other nodes and are predicted to have the greatest impact on the flow of information throughout the network.

Hub nodes can be identified by measurements of the centrality of a network which themselves are calculated by analyzing the overall topology of the network. To date, more than 100 centrality measures have been identified during the assessment of different network nodes and several tools, plugins and packages have been developed for the calculation of these measures (Jalili et al, 2015) (Barabási & Oltvai, 2004). Furthermore, some tools have been developed for the identification of hub nodes of a network based on its centrality measures. The simplest local centrality measure of a graph is the degree centrality. Betweenness centrality is a global centrality measure and one of the most important parameters for the identification of network hubs (Freeman et al, 1991). Betweenness is defined as the tendency of a node to be on the shortest path between nodes in a graph (Frainay & Jourdan, 2017). Nodes with high betweenness are considered as influencers of information flow within a network (Oldham et al, 2019). Neighborhood connectivity is a (semi-) local centrality measure of a network that deals with the connectivity (number of neighbors) of nodes. We call it a semi-local metric as it is not restricted to only first neighbors of a node and encompasses a broader environment. The neighborhood connectivity of a node is defined as the average connectivity of all neighbors of that node (Maslov & Sneppen, 2002). It is also reported that not only the number of first connections of a node (degree centrality), but also the extent to which the immediate neighbors of the node are connected with each other and other nodes (neighborhood connectivity) is determinant of the importance of a node in the network (Bhola et al). Other network metrics such as network density (Farrar et al, 2018), size (Farrar et al, 2018), path length (Blain-Moraes et al, 2017), PageRank versatility (Gao et al, 2017) and modularity (Blain-Moraes et al, 2017) can further be used to evaluate the topology of networks. Among all these measures, degree and betweenness centrality are of the most commonly used metrics for the identification of network hubs, especially in biological contexts.

The simultaneous consideration of a set of centrality measures has been previously used as a strategy to find the most influential network nodes. del Rio et al. (2009) by analyzing 16 well-known centrality measures demonstrated that while identification of network hubs based on a single centrality measure is not statistically significant, the combination of two centrality measures that involve both local and global features of the network could more reliably predict the most influential nodes (del Rio et al, 2009). In systems biology studies, nodes with high degree and betweenness centrality are usually represented as network hubs in the context of defining both local significance and global network flow (Fu et al, 2015; Oulas et al, 2019). However, no method or algorithm has been developed to date that integrates these centrality measures with the purpose of synergizing their effects. Additionally, in many networks there are nodes that are topologically positioned in the center of the network and consequently have a high degree centrality but low betweenness centrality due to their lack of connections to nodes outside the main module (Oldham et al, 2019). Specifically, nodes may exhibit a high local centrality but low global centrality, or vice-versa, depending on their position in the network (Guimerà & Nunes Amaral, 2005). Betweenness centrality measurements are therefore biased by their position in the network and consequently should be carefully used for identification of network hubs or in developing new hub identification algorithms. Though, the positional bias of betweenness centrality has not been clearly addressed in previous studies and no solution has been proposed to correct for this computationally.

To overcome these problems, we developed a novel formula we termed the integrated hubness score (IHS) that integrates the most significant network centrality measures in order to synergize their effects and simultaneously remove their biases to identify the most essential regulatory molecules in a network. The IHS is the first method that truly integrates the effect of degree and betweenness centrality. To address the issue of positional bias of betweenness centrality we utilized one of the other network centrality measures named neighborhood connectivity. Considering all centrality metrics represented in the literature, we believe that three metrics including degree centrality, neighborhood connectivity, and betweenness centrality are most important for the identification of network hubs in an integrative manner. Each of these centrality measures captures a different topological dimension of the graph including local, semi-local, and global topology. One of the other advantages of these centrality measures is that none of them require a fully connected graph or module to be calculated. In this study, we addressed the problems and gaps in integration of centrality measures and identification of true network hubs. We first precisely interrogated the association of neighborhood connectivity with betweenness centrality and determined that neighborhood connectivity is an effective method for removing positional bias in the context of betweenness centrality measurements. Next, we compared the IHS method with current methods of hub identification. Overall, our results reveal that the IHS algorithm out performs all methods tested in the identification of the most influential network nodes.

## Results

We aimed to integrate the most commonly used local and global measures of network centrality, namely degree centrality and betweenness centrality respectively, and to synergize their effect for the identification of influential nodes in the network in an unbiased way. However, betweenness centrality is biased by its position in the network and the edge numbers of its surrounding local environment. By contrast, neighborhood connectivity represents the average number of edges connected to immediate neighbors and consequently is a good candidate metric for removing the positional bias introduced by betweenness centrality measures. As an example, the topological analysis of a PPI network of adrenocortical carcinoma (ACC), clearly depicted that there is a positional bias in the distribution of betweenness centrality scores between nodes in a network which is contrary to neighborhood connectivity distribution (Figure 1). Accordingly, we first inspected the association of betweenness centrality and neighborhood connectivity from different aspects including linearity, monotonicity, and dependence which altogether reveal the nature of the association of two variables.

**Figure 1.**
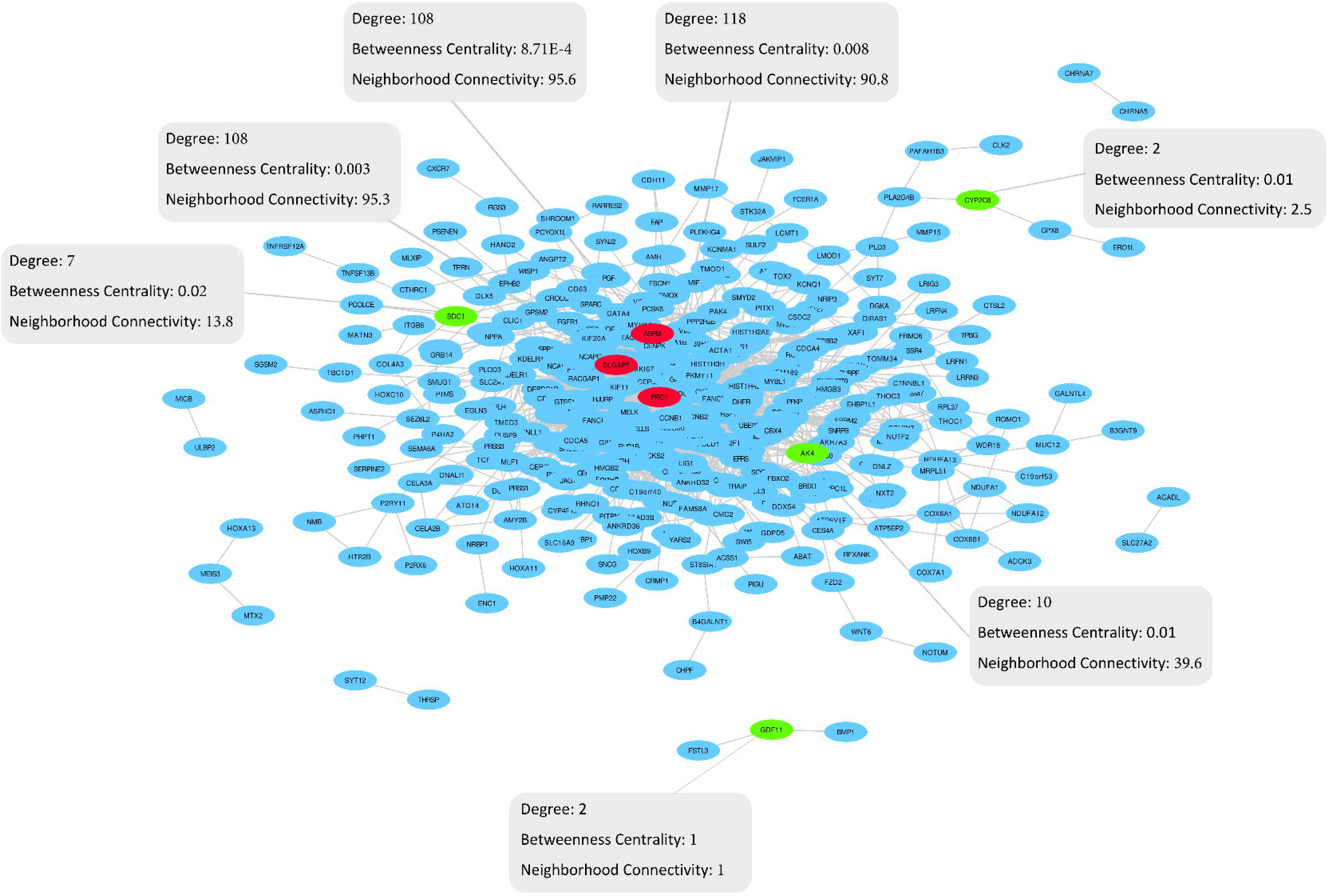
The protein-protein interaction network of adrenocortical carcinoma. This network has been reconstructed using the interactions presented by the STRING database and based on genes in the blue module provided by (Xia et al, 2019) according to the weighted gene co-expression network analysis (WGCNA) as seed proteins. Red nodes are three examples of nodes with high neighborhood connectivity and low betweenness centrality in the center of the network; while, green nodes are representative of nodes with lower neighborhood connectivity and higher betweenness centrality on the edges of the network. The network was illustrated by Cytoscape and analyzed by the NetworkAanalyzer.

### Betweenness centrality is dependent on neighborhood connectivity

In order to examine if betweenness centrality and neighborhood connectivity are correlated with each other and to investigate the nature of their association we built a computational pipeline through which the innate features of these metrics and their dependence were carefully assessed. The normality assessments demonstrated that betweenness centrality and neighborhood connectivity of nodes are non-normally distributed (*p*-value ≅ 0) in all networks, and the majority of studied networks (Figure 3-a), respectively (Figure 3-b). Also, non-linearity/monotonicity assessments indicated that these two centrality measures are non-linearly/non-monotonically correlated with each other (estimated degree of freedom of smooth terms > 1, *p*-value < 0.05) in both of the independent real-world biological networks analyzed. Similarly, non-monotonicity evaluations revealed that the selected centrality measures were more non-monotonically correlated (Multiple R-squared from rank-regression > squared Spearman’s rank correlation coefficient) with each other rather than monotonically. Also, CANOVA and Hoeffding’S independence analyses indicated that betweenness centrality is significantly dependent on and non-linearly/non-monotonically correlated with neighborhood connectivity (*p*-value < 0.05; Figure 3-c). Moreover, both non-linear non-parametric statistics (NNS) descriptive dependence and Hoeffding’S independence analyses demonstrated that betweenness centrality is dependent on neighborhood connectivity (Figure 3-d).

**Figure 2.**
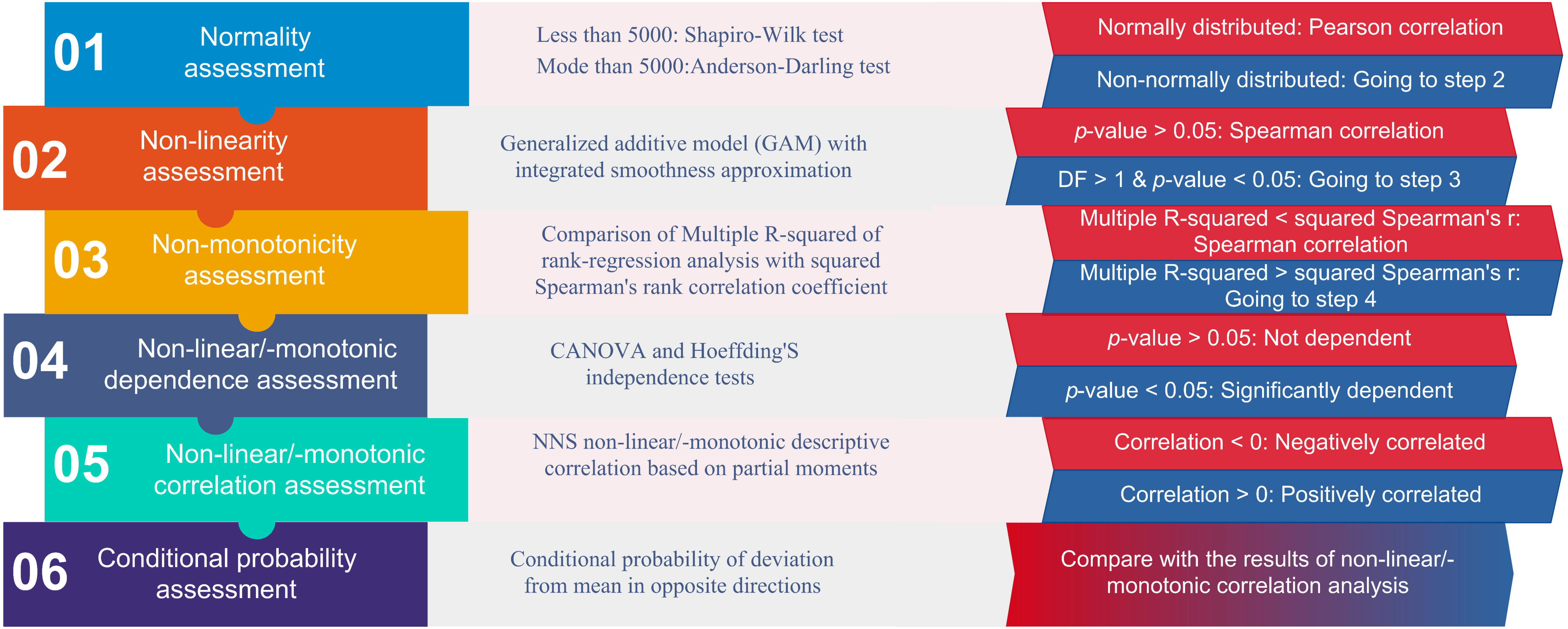
The assessment workflow of the association of two network centrality measures. This workflow illustrates the step-wise assessment of innate characteristics and association/dependence of two centrality metrics of a network. The extended form of abbreviations used in the figure are as follows; DF: degree of freedom, CANOVA: continuous analysis of variance, NNS: non-linear non-parametric statistic.

**Figure 3.**
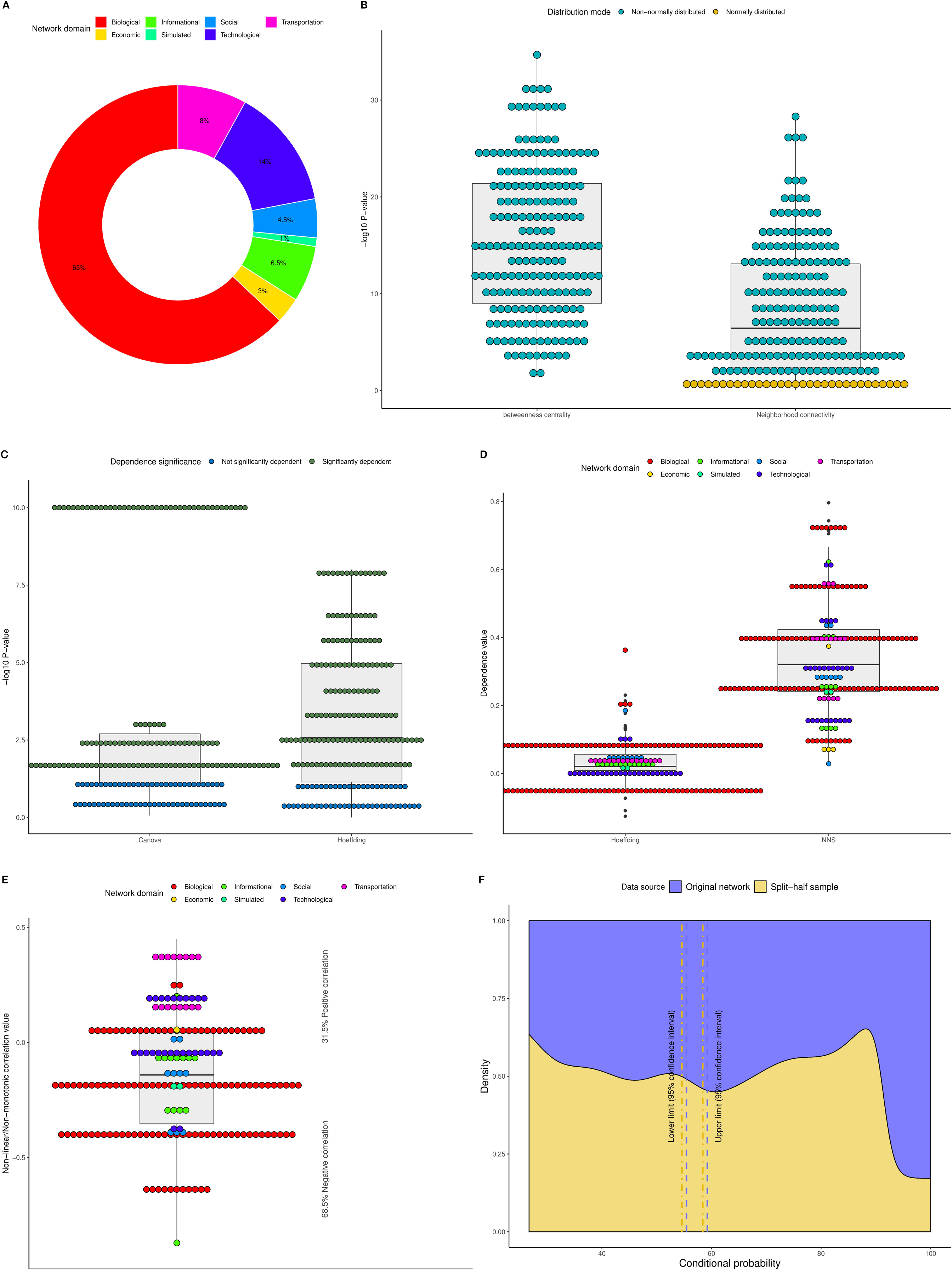
Distribution mode and association of betweenness centrality and neighborhood connectivity in 200 networks. **A**. The proportion of network domains among 200 networks. **B**. Distribution mode (i.e. Gaussian or non-Gaussian distribution) of the selected centrality measures across all under-investigation networks. **C**. Statistical significance of the dependence of betweenness centrality to neighborhood connectivity based on two different dependence tests including continuous analysis of variance (CANOVA) and Hoeffding across all networks. **D**. Descriptive dependence of betweenness centrality to neighborhood connectivity based on two different tests including non-linear non-parametric statistic (NNS) and Hoeffding across all networks. **E**. Non−linear/Non−monotonic descriptive correlation of betweenness centrality and neighborhood connectivity across all networks. **F**. The conditional probability of deviation of betweenness centrality from its mean given that neighborhood connectivity has deviated from its corresponding mean in the opposite direction based on both original networks as well as their split-half random samples.

### Neighborhood connectivity is the right choice for unbiasing betweenness centrality

In order to further test if neighbourhood connectivity is a good candidate for removing the positional bias of betweenness centrality we did partial moment-based correlation analyses followed by conditional probability assessments. The NNS descriptive correlation analyses based on higher-order partial moment matrices indicated that betweenness centrality and neighborhood connectivity are negatively non-linearly/non-monotonically correlated with each other in a significant proportion of networks including the majority of biological networks analysed except in transportation and economic domains (Figure 3-e). In addition to correlation and dependence assessment, we performed a conditional probability analysis as a complementary test to further examine the association of under-investigation centrality measures. The conditional probability measurements based on both whole networks, as well as their split-half random samples, determined that betweenness centrality and neighborhood connectivity of a significant fraction of nodes deviate from their corresponding means in opposite directions (Figure 3-f). Furthermore, analysis of innate characteristics and association of betweenness centrality and neighborhood connectivity of all nodes of all 200 under-investigation networks together demonstrated consistently that betweenness centrality is dependent on and negatively correlated with neighbourhood connectivity. Also, for the majority of nodes of all 200 networks, these centrality measures deviate from their means in opposite directions (Table 1).

**Table 1.**
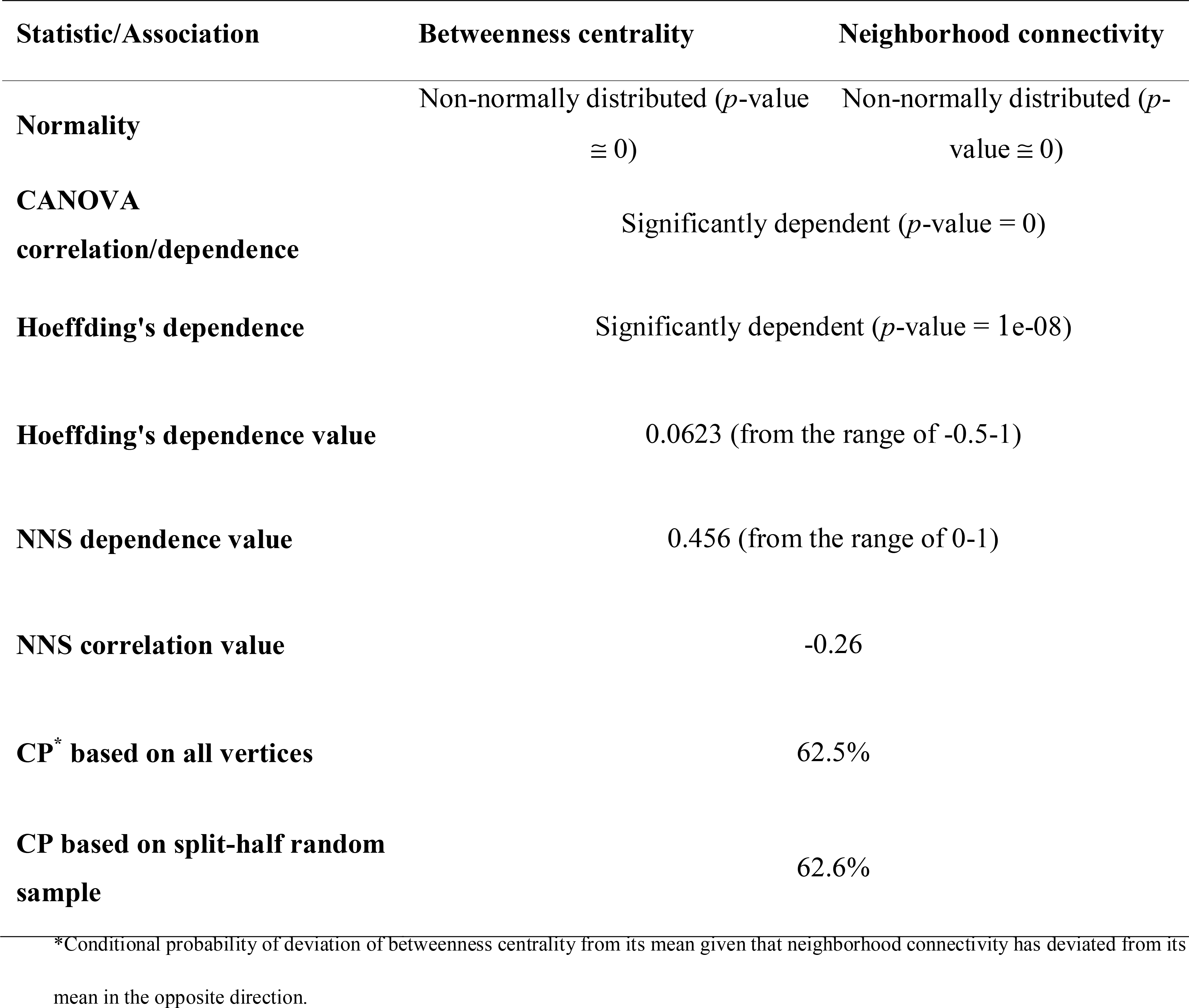
Innate characteristics and association of betweenness centrality and neighborhood connectivity of entire vertices of all 200 networks.

### IHS formula

In order to integrate degree and betweenness centrality and calculate their synergistic effect the multiplication method was used. However, as explained above, betweenness
centrality of each node has a positional bias in the network and consequently multiplication of betweenness centrality by degree results in a positional biased synergistic product. Thus, betweenness centrality should be normalized in a way that does not disrupt its ratio among network nodes. Accordingly, the positional bias of betweenness centrality was removed via multiplication by neighborhood connectivity owing to the following results. Based on the non-linear/non-monotonic (higher-order partial moment based) correlation and dependence analyses, betweenness centrality had a negative non-linear correlation with neighborhood connectivity in the majority of networks studied. Also, the conditional probability assessments indicated that betweenness centrality and neighborhood connectivity of a significant proportion of nodes in all networks deviated from their corresponding means in opposite directions. However, as the results of multiplication of betweenness centrality by neighborhood connectivity were still skewed, they were log2 transformed to further stabilize their variance and bring them closer to an unbiased distribution across orders of magnitude. In the end, the result, namely unbiased betweenness centrality, was multiplied by degree centrality and the IHS was calculated.

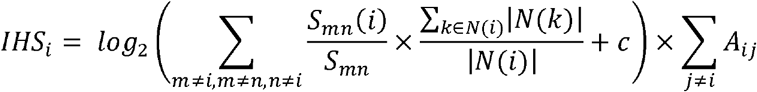

Where N and E are representative of neighbors and edges respectively, *S*_*mn*_ is the number of shortest paths between nodes *m* and *n*, *S*_*mn*_ (i) is the number of shortest paths between nodes *m* and *n* which pass through node *i*, N (i) is set of neighbors of node *i*, and *c*, is the constant.

Or, in short

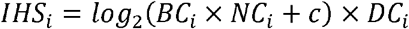

In other words, IHS is the synergistic product of the most important pure local centrality measure (i.e. degree centrality), an impactful semi-local centrality measure (i.e. neighborhood connectivity) and the most influential global centrality metric (i.e. betweenness centrality) in a way that simultaneously removes the positional bias of betweenness centrality.

### IHS outperforms other hub identification methods in detecting influential network nodes

The comparison of IHS with four other hub identification methods including corresponding paper represented hubs, degree centrality, Kleinberg’s hub centrality scores (Kleinberg, 1999) and maximal clique centrality (MCC (Chin et al, 2014)) in practice demonstrated that IHS generally outperformed other methods. In other words, according to the literature-based text mining through the title and abstract of all available articles within the PubMed database, hubs identified by the IHS method had the highest number of reported associations with the context of two out of four under-investigation corresponding studies. Also, the IHS-based identified hubs were ranked second and third in two other studies in comparison to all four other methods. That is to say, altogether, the hub nodes identified by the IHS method had the greatest number of reported associations with the context of their corresponding studies (Figure 4 and Table EV2).

**Figure 4.**
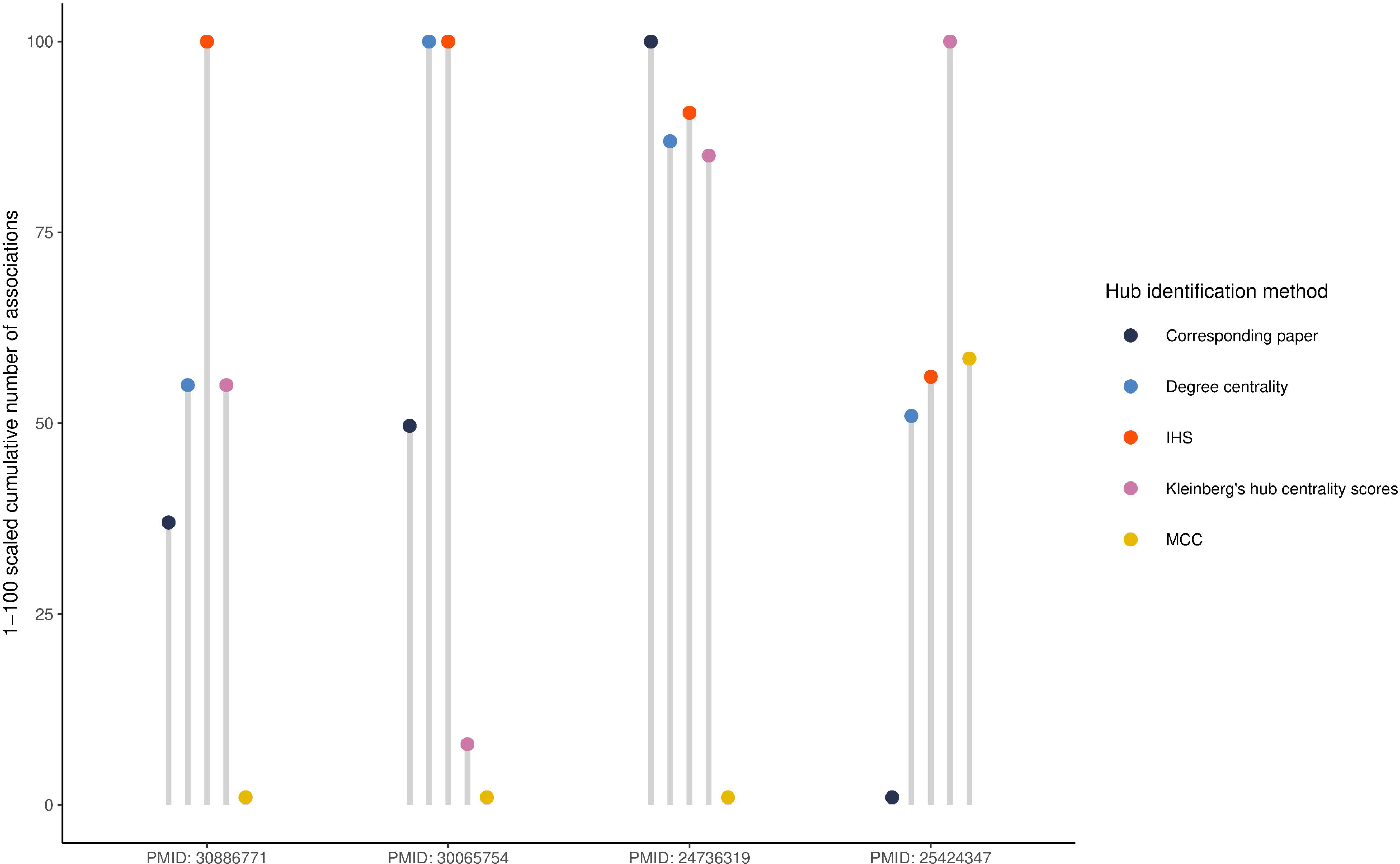
Comparison of IHS with four other hub identification methods in practice. The graph illustrates 1-100 normalized cumulative number of associations of identified hubs by each of hub identification methods with the context of their respective studies according to the title and abstract screening of all available literature on the PubMed database. The “Corresponding paper” refers to the network hubs suggested by the respective paper itself. IHS and MCC are abbreviations of the integrated hubness score and maximal clique centrality, respectively.

## Discussion

Identification of the most influential nodes is a necessity in all network analyses across all fields, and different centrality measures are being used for this purpose. Several methods and algorithms have also been proposed for the identification of network hubs during previous decades. In recent years, most of the biological studies use the weighted gene co-expression network analysis (WGCNA; (Langfelder & Horvath, 2008)) method for identification of functional modules and degree-based hub nodes (Liu et al, 2019). Similarly, the CINNA method first identifies the giant component of the network and subsequently suggests some centrality measures based on PCA for the selection of network hubs (Ashtiani et al, 2019; Ashtiani et al, 2018). However, these methods would lose a subclass of network hubs, namely inter-modular hubs (Ferreira et al, 2013; Missiuro et al, 2009), which are nodes that could play an influential role in the flow of information or function between different functional modules in complex networks. Thus, integration of the most important centrality measures that capture all topological dimensions of a network and synergizing their impacts could be a big step towards identification of the most influential nodes. Furthermore, some centrality measures including betweenness centrality, the most important global centrality metric, are biased by their positions (the edge numbers of their surrounding local environment) in the network. Freeman has proposed a formula for the normalization of betweenness centrality (Freeman, 1978); however, this formula adjusts the betweenness centrality for the network size, not its positional bias. This issue is also addressed in the IHS formula. On one hand, neighborhood connectivity is highly representative of the size of surrounding local environment. On the other hand, according to the methodology applied for the assessment of dependence and correlation of betweenness centrality and neighborhood connectivity, betweenness centrality is dependent on and negatively correlated with neighborhood connectivity. Therefore, neighborhood connectivity is the best centrality measure for removing the positional bias of betweenness centrality and consequently was used in the IHS formula.

The IHS, while being very simple and fast to be calculated, is an unsupervised method that generates the synergistic product of the most important local, semi-local, and global centrality measures in a way that simultaneously removes the positional bias of betweenness centrality for the identification of hub nodes in the whole network as well as its functional modules. Moreover, the IHS is not dependent on the arbitrary threshold selection. On the contrary, a list of most influential nodes (hubs) of a particular network can be identified by sorting the set of nodes based on their IHS value. The required centrality measures and the IHS of each node could be calculated using the “influential” R package (link). It is also noteworthy that using the “influential” R package, neighborhood connectivity is for the first time calculable in the R environment. Additionally, the centrality measures calculated by other tools such as Cytoscape software could be imported into the R environment for the calculation of IHS of nodes.

The comparison of IHS formula with four other hub identification methods in identification of true network hubs confirmed that the IHS method outperforms these algorithms. Precisely speaking, according to a text mining based on the title and abstract of the literature within the PubMed database, hubs identified by the IHS method had the highest number of reported associations with the context of two out of four under-investigation corresponding studies and were ranked second and third in two other ones. Altogether, therefore, the hub nodes identified by the IHS method had the greatest number of reported associations with the subjects of the studies, which confirms the power of the IHS method in identification of the most influential network nodes. Although simultaneous existence of two terms (i.e. the identified hub node and the context of the research) in the title and/or abstract of the literature does not necessarily imply causality, but it could be informative of an association, no matter direct or indirect, between the hub node and the context of the study. It is also worth mentioning that some of the hub nodes might contain driver genes of the disease under investigation but due to their novelty, no previous association has been reported. For example, in the ACC gene network, the association of four out of seven hub genes that were specifically identified by the IHS method but not represented by the corresponding paper including AURKB (Borges et al, 2013; Mohan et al, 2019; Subramanian & Cohen, 2019), CDK1 (Nilubol et al, 2018), NDC80 (Xiao et al, 2018) and PLK1 (Bussey et al, 2016) with ACC have been previously validated by biological assays. Though no biological experiment has been previously done regarding the association of three other hub genes with the ACC, their high IHS is suggesting that they might play pivotal roles in the carcinogenesis of ACC and should be considered in the future studies. Similarly, in the comparison of IHS-based hubs and the corresponding paper reported network hubs of the other three studies, the association of the majority of IHS-based uniquely identified hub genes with the disease under-investigation have been previously experimentally validated.

In conclusion, the IHS method we describe here is based on the accurate evaluation of the association of most important network centrality measures in order to integrate them in such a way that their powers are synergized and the positional biases are removed. Also, the workflow depicted in Figure 2 could be used as a reference for the assessment of association and dependence of network centrality measures as well as any other two continuous variables. According to the results, the IHS formula outperforms other algorithms in identifying the most influential nodes in terms of both accuracy and specificity. Moreover, as the IHS is an unbiased synergistic product, it is very general and is not dependent on either directedness or weightedness of the network and can be calculated for directed and weighted networks as well. Additionally, the IHS method is applicable to both the statistically inferred networks as well as the experimental real-world ones. All in all, as the IHS is an unbiased synergistic product of most important centrality measures that altogether cover all topological dimensions of the network and according to our results, we believe that the IHS could identify true hubs and the most influential nodes in a network which could be a great benefit for all future systems biology studies.

## Materials and Methods

### Reagents and Tools table

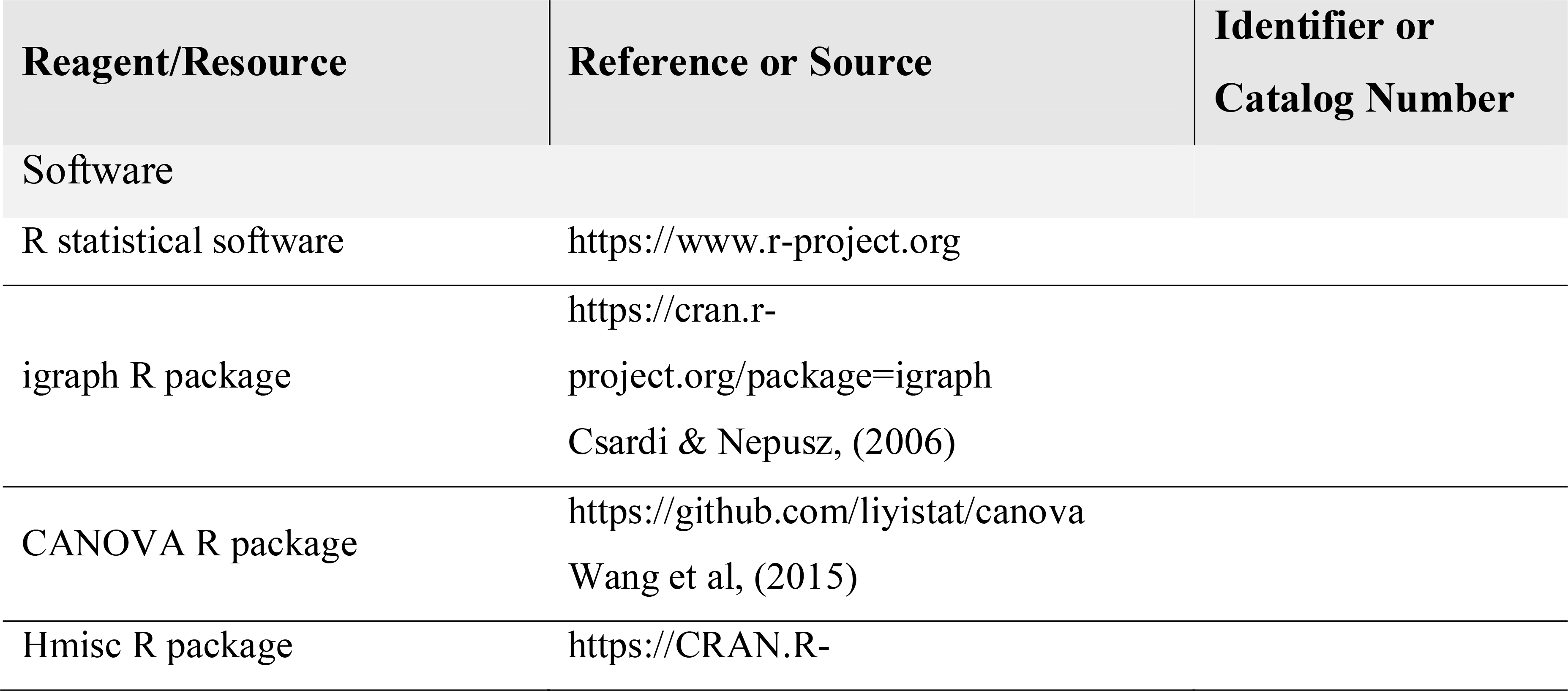
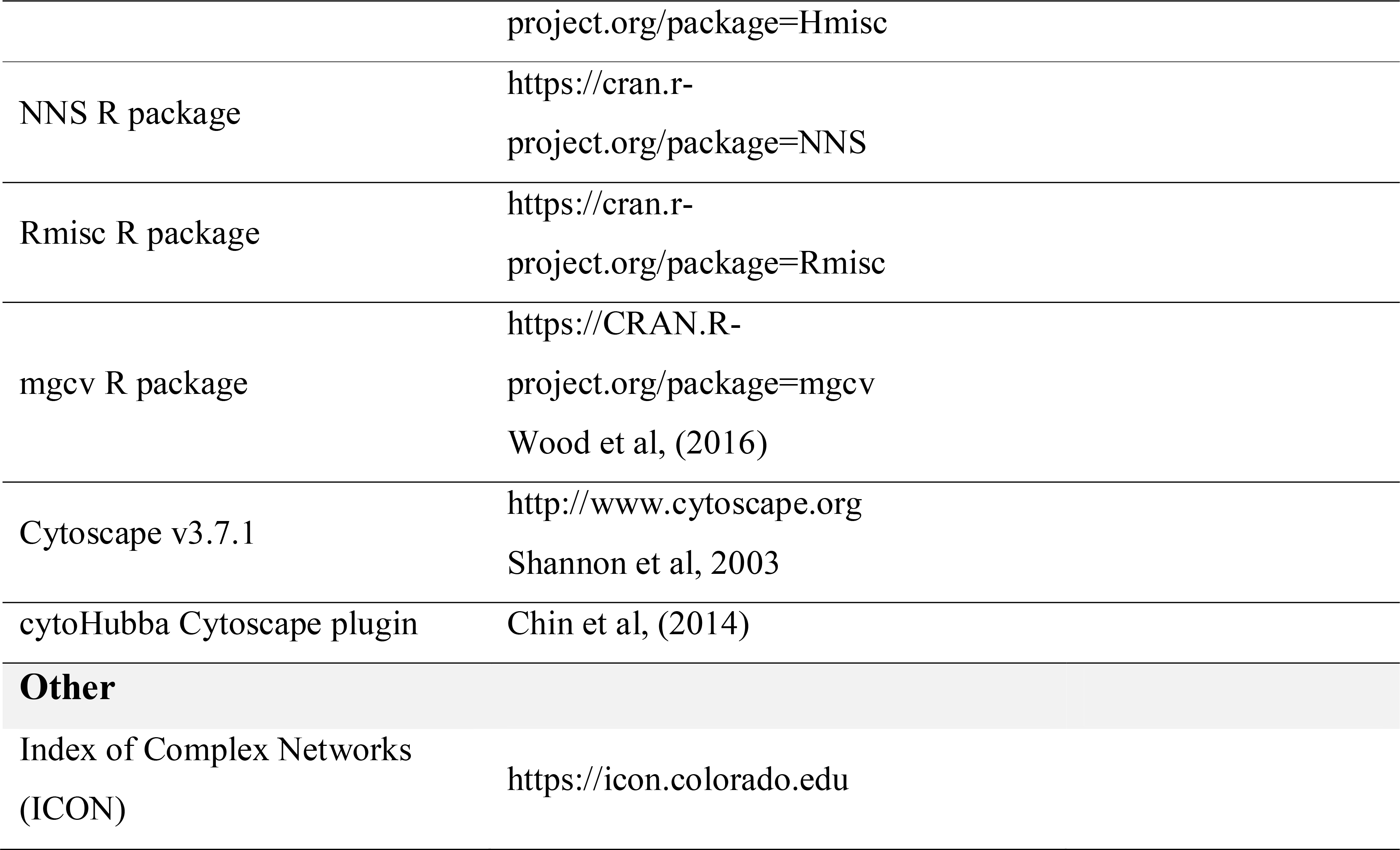

| Reagent/Resource | Reference or Source | Identifier or Catalog Number |
| --- | --- | --- |
| Software |  |  |
| R statistical software | https://www.r-project.org |  |
| igraph R package | https://cran.r-project.org/package=igraph<br>Csardi & Nepusz, (2006) |  |
| CANOVA R package | https://github.com/liyistat/canova<br>Wang et al, (2015) |  |
| Hmisc R package | https://CRAN.R- |  |

|  |  |
| --- | --- |
|  | [project.org/package=Hmisc](https://cran.r-project.org/package=Hmisc) |
| NNS R package | https://cran.r-project.org/package=NNS |
| Rmisc R package | https://cran.r-project.org/package=Rmisc |
| mgcv R package | https://CRAN.R-project.org/package=mgcv<br>Wood et al, (2016) |
| Cytoscape v3.7.1 | http://www.cytoscape.org<br>Shannon et al, 2003 |
| cytoHubba Cytoscape plugin | Chin et al, (2014) |
| **Other** |  |
| Index of Complex Networks (ICON) | https://icon.colorado.edu |

### Methods and Protocols

#### Data preparation

A total of 200 connectivity tables including 198 real adjacency matrices as well as 2 random ones generated via igraph R package (Csardi & Nepusz, 2006) were gathered for the analysis of the topology of networks and the association of their selected centrality measures. The real-world networks included 196 adjacency matrices compiled by Ghasemian et al. (Ghasemian et al, 2019) from the Index of Complex Networks (ICON) (https://icon.colorado.edu) and two of them downloaded from independent biological studies which included a PPI from (Xia et al, 2019) and a miRNA expression dataset from (Yepes et al, 2016). The connectivity matrices were retrieved from different domains to ensure the reliability of our analyses. A list of all networks is provided in Table EV1. Additionally, a PPI dataset, as well as its associated co-expression dataset, from (Xia et al, 2019) and three PPI networks from (Liu et al, 2018),(Zhang et al, 2014), and (Nair et al, 2014) were obtained from the corresponding papers to compare and assess the applicability of our method in comparison with other available methods in identification of nodes with the highest impact in the network considering the context of the study of each paper.

#### Network reconstruction and analysis

A correlation analysis was done based on the Pearson algorithm for the identification of co-expressed genes in miRNA and protein expression datasets independently. Next, an undirected network was reconstructed for each of these co-expression/PPI datasets, all other real-world connectivity matrices retrieved from ICON as well as the random adjacency matrices via the igraph R package. Then, the topology of each network and its centrality measures were analyzed using the pre-built functions of the igraph R package as well as a novel R function for the calculation of neighborhood connectivity. The Cytoscape software v3.7.1 was also used for network visualization purposes (Shannon et al, 2003). All of the downstream analyses and assessments were done both independently for each network as well as across all networks as a whole.

#### Assessment of the association of neighborhood connectivity and betweenness centrality

From a statistical viewpoint, the intrinsic features of variables should be inspected prior to association analyses. Similar to real-world networks that their parts are interdependent and follow non-linear associations (De Domenico et al, 2014; Jiang & Lai, 2019), their centrality measures might also be non-linearly/non-monotonically correlated to each other. Thus, though previously done by other researchers (Li et al, 2015; Oldham et al, 2019), ranking-based monotonic correlation tests such as Spearman’s rank correlation test does not produce correct and reliable enough results. On the other hand, assessment of local associations of two continuous variables with a subsequent global assessment of all local correlations would more reliably and correctly assess non-linear non-monotonic correlations. However, the correlation between two variables is not enough for proving causality. While correlation analysis could indicate a desired predictive relationship, dependence analysis, which is one of the sub-branches of correlation tests, could reveal the statistical relationship between two variables (Wang et al, 2015). Also, conditional probability assessment has been proposed as a complementary test to dependence analysis for the investigation of causality (van Rooij & Schulz, 2019). Accordingly, we considered assessing dependence, correlation, and the conditional probability of opposite behaviors of neighborhood connectivity and betweenness centrality in order to reach more reliable conclusions.

First, Gaussian distribution of selected centrality measures, namely betweenness centrality and neighborhood connectivity, was assessed using Shapiro–Wilk test or Anderson-Darling test for variables with less than or more than 5000 objects, respectively. Next, the statistical significance of dependence of betweenness centrality to neighborhood connectivity and the non-linear/non-monotonic correlation between them was assessed by two methods including continuous analysis of variance (CANOVA) and Hoeffding’S independence tests using CANOVA (Wang et al, 2015) and Hmisc (https://CRAN.R-project.org/package=Hmisc) R packages, respectively. CANOVA test is able to detect dependence and non-linear/non-monotonic correlation between two continuous variables (Wubetie, 2019). Furthermore, CANOVA test works well and is robust in non-linear correlation cases, especially when the association between two continuous variables is non-monotonic (Wang et al, 2015). Subsequently, based on the non-linear non-parametric statistics (NNS), descriptive correlation and dependence of the selected centrality measures were analyzed using the NNS R package (https://cran.r-project.org/package=NNS). The NNS is a robust method for the assessment of dependence and correlation of two variables with non-linear/non-monotonic association and uses higher-order partial moment matrices instead of global measurements. In other words, the NNS method calculates the correlation coefficient by combining linear segments resulted from ordered partitions without the need to perform a linear transformation. Also, as a complementary test, the conditional probability of deviation of betweenness centrality and neighborhood connectivity from their corresponding means in the opposite direction was calculated in each network as well as across all networks. Additionally, the split-half random sampling method was used for each network as well as all networks as a whole for reliability assessment of conditional probability assessments. At last, the 95 percent confidence interval of all conditional probability assessments was calculated using the Rmisc R package (https://cran.r-project.org/package=Rmisc).

#### Interrogation of the non-monotonic association of neighborhood connectivity and betweenness centrality

Though a single regression line does not fit all models with a certain degree of freedom and consequently is not applicable in high-throughput assessments, regression analysis was used to more precisely evaluate non-linearity and non-monotonicity of the association of betweenness centrality and neighborhood connectivity of 2 real-world independent biological networks. For this purpose, the non-linear correlation between neighborhood connectivity and betweenness centrality was interrogated by fitting a generalized additive model (GAM), with integrated smoothness approximation using the mgcv R package (Wood et al, 2016), which estimates nonparametric functions of the predictor (independent) variable, namely neighborhood connectivity. GAM is a technique for regression analysis of non-linear/non-monotonic associations (Faraji Gavgani et al, 2018). Subsequently, the most squares strategy was used to decipher if the association of selected centrality measures is more of a monotonic or a non-monotonic form. Accordingly, squared coefficients of the correlation of betweenness centrality and neighborhood connectivity based on Spearman’s rank correlation analysis and ranked regression test with non-linear splines were compared, and the correlation was designated as monotonic if squared coefficient of Spearman’s rank correlation was higher compared with the other test and was identified as non-monotonic if the argument was inverse. For the ranked regression analysis, the splines R package was used to generate a basis matrix for natural cubic splines of the predictor variable, namely neighborhood connectivity. A schematic workflow of the methods implemented for the assessment of innate features and association of selected network centrality measures is shown in Figure 2.

#### Evaluation of the performance of IHS formula in comparison with other methods

To evaluate the functioning of IHS formula in real-world biological networks three different studies on ACC, gastric cancer, hepatocellular carcinoma, and coronary artery disease were investigated. To this end, firstly, hub nodes were identified based on five different methods including corresponding paper represented hubs, degree centrality, Kleinberg’s hub centrality scores, MCC (the recommended method by authors of cytoHubba plugin (Chin et al, 2014)), as well as the IHS formula. Degree centrality and Kleinberg’s hub centrality scores were calculated by the igraph R package, and MCC was calculated by the cytoHubba plugin of Cytoscape software. It is also worth mentioning that the same number of top nodes used in each paper was applied for the selection of hub nodes by all other methods. Then, the association of identified hubs with the context of each respective study was inspected based on screening of title and abstract of all available papers on the PubMed database (www.ncbi.nlm.nih.gov/pubmed/; December 5-10, 2019) and the cumulative number of returned items considering all hubs was calculated for each method. Finally, the cumulative number of the results of each hub identification method was compared with that of the IHS method.

## Data availability

All datasets generated/analyzed for this study are included in the manuscript and the Expanded View files. The computer code produced in this study are available in the following databases:

- The “influential” R package for the calculation of IHS and its required centrality metrics: CRAN (https://cran.r-project.org/package=influential)

## Acknowledgments

The authors would like to thank two biostatisticians, Parinaz Mehdipour and Ehsan Rezaei-Darzi, for their consultations regarding proper use of statistical methods and evaluations. Also, the constructive feedbacks on the manuscript by Hieu Tri Nim are highly acknowledged. The results shown in this study are in part based upon data generated by the TCGA Research Network: http://cancergenome.nih.gov/. This work was supported by Monash University.

## Author contributions

AS, MR and PC conceptualized the study, AS performed data analysis and writing of the manuscript, with contributions from MR and PC.

